# A two-oscillator SCN model with period adaptation and systemic feedback captures photoperiod and T-cycle aftereffects *in vivo* and in explants

**DOI:** 10.64898/2026.08.09.743784

**Authors:** Vuong Hung Truong, Jihwan Myung

## Abstract

Light history leaves persistent changes in circadian period, but where this history is stored remains unresolved. Suprachiasmatic nucleus (SCN) network models have often approached photoperiodic encoding through phase organization or coupling strength. We computationally tested slow adaptation of subregion-specific intrinsic periods as an alternative memory mechanism. The model asymmetrically couples dorsal (*D*) and ventral (*V*) SCN oscillators and adds a systemic oscillator (*X*) representing putative circadian feedback present *in vivo* but lost *ex vivo*. With a single parameter set, period adaptation captured the direction and approximate magnitude of behavioral aftereffects across photoperiod and T-cycle conditions. Adapting coupling strength instead of period failed to reproduce the *V*-leading-*D* phase order reported after T22. Removing systemic feedback preserved the photoperiod-dependent period ordering but inverted the T22 and T26 aftereffects, an inversion that matched SCN explant observations. The model also yielded distinct *D*-*V* phase organization for each of 18:6 LD, T23, and T25. These results suggest that subregion-specific period plasticity provides a parsimonious substrate for encoding light history, while the dependence on systemic feedback indicates that behavioral period may not be a readout of the SCN alone.

**Graphical Abstract:** 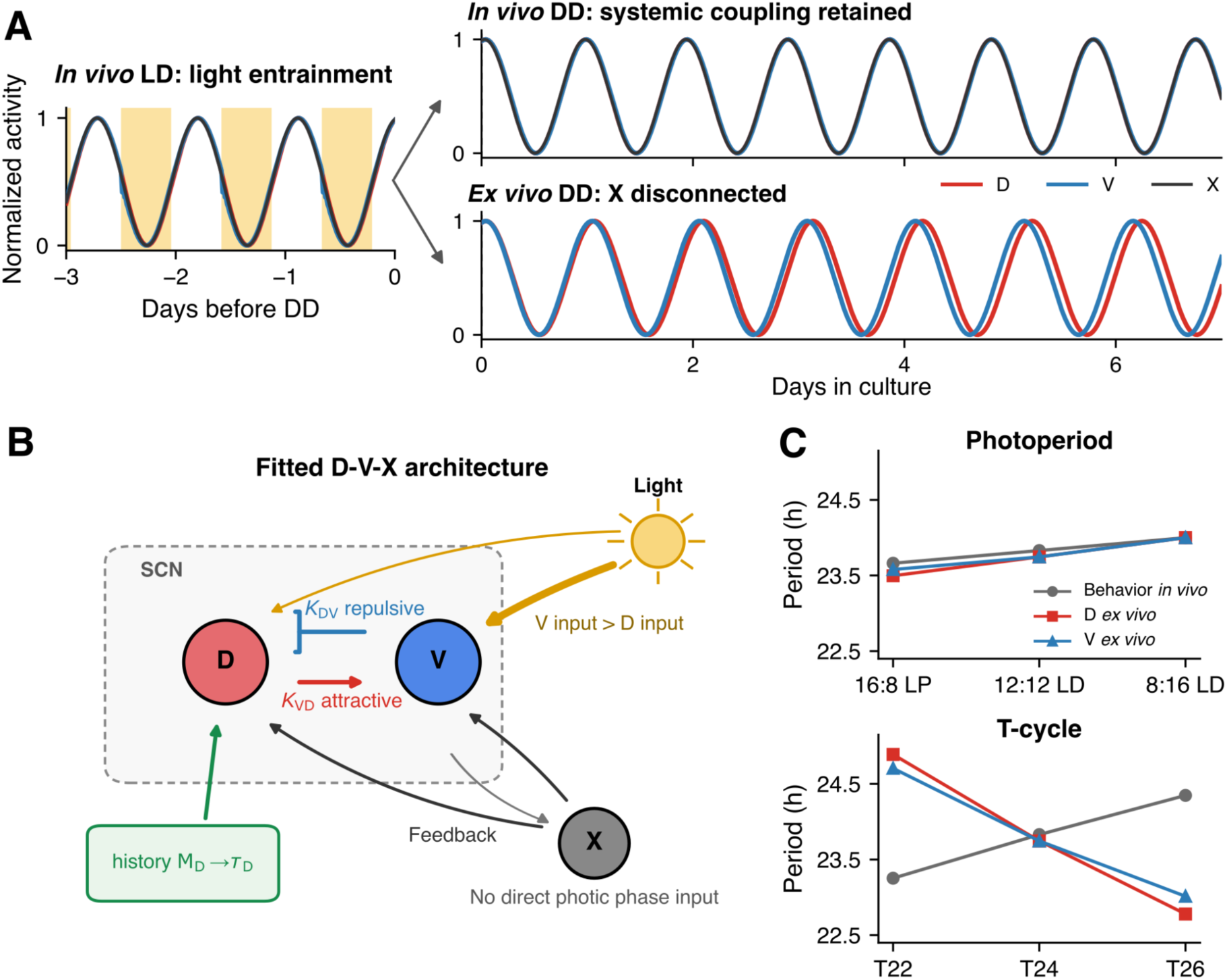

**A model with dorsal period adaptation and phenomenological systemic feedback accounts for behavioral and explanted SCN aftereffects. (A)** During T22 entrainment, dorsal (*D*), ventral (*V*), and systemic (*X*) oscillators remain phase-locked. After release into constant darkness, systemic coupling maintains a unified *in vivo* rhythm, whereas removing *X* feedback in the explant simulation allows the *D*-*V* network to express a distinct *ex vivo* period aftereffect. (B) The SCN model is modeled as an asymmetrically coupled attractive-repulsive oscillator network with stronger photic input to *V*. Light history is encoded via plasticity of the intrinsic period in *D*, while *X* represents putative systemic circadian feedback available *in vivo* and lacking direct photic input. (C) The model reproduces concordant period changes in behavior and SCN explants across photoperiods, but opposing period changes following T-cycle entrainment.

## Introduction

Circadian rhythms are endogenously generated cycles of approximately 24 h in physiology and behavior (Vitaterna et al., 2001). Underlying these rhythms is the molecular clockwork, composed of interlocked transcriptional-translational feedback loops that are entrained by environmental time cues (zeitgebers), most notably the light-dark cycle. Through entrainment, organisms align their internal physiology with the external world, optimizing energy use, cognitive performance, and survival (Patke et al., 2020).

In mammals, the central circadian pacemaker is located in the suprachiasmatic nucleus (SCN) of the hypothalamus. The SCN tracks the external light-dark cycle and coordinates rhythms in the brain and peripheral tissues (Moore et al., 2002). It receives direct photic input from intrinsically photosensitive retinal ganglion cells (ipRGCs), whose axons form the retinohypothalamic tract (RHT) and make synaptic contacts in both dorsal (*D*) and ventral (*V*) subregions (Fernandez et al., 2016), although the activation threshold appears higher in *D* than in *V* (Starnes & Jones, 2023). Individual variation in circadian behavior, whether due to age or genotype, is often linked to changes in SCN properties. For instance, age-related shortening of the behavioral circadian period in rats is paralleled by a shortened intrinsic period of *Per1-luc* rhythms in SCN explants (Yamazaki et al., 2002). The period of SCN firing rate rhythms, both *in vivo* and *ex vivo*, closely mirrors the behavioral period determined by genotype (Albus et al., 2002; Herzog et al., 1998; Honma et al., 1998; Liu et al., 1997; Nakamura et al., 2002; Welsh et al., 1995; Yamazaki et al., 2002). The causal role of the SCN in determining circadian period is further supported by transplantation studies. Grafting SCN tissue from *tau*-mutant hamsters into SCN-lesioned, arrhythmic hosts was sufficient to restore behavioral rhythmicity with a period determined by the donor (Ralph et al., 1990).

While the SCN keeps time with remarkable stability and precision under unchanging conditions of 12:12 light-dark (LD) or constant darkness, it also displays a degree of plasticity by adaptively shifting its dynamics in response to changes in light schedules. Anatomically and functionally, the SCN is frequently described in terms of a dorsal-ventral (shell-core) organization, a coarse-grained description of a heterogeneous and partly overlapping neuronal network. This broad organization is widely conserved in mammals and delineated by neuropeptide expression: the *V* subregion is enriched in vasoactive intestinal peptide (VIP)-expressing neurons and responds rapidly to light, and the *D* subregion is enriched in arginine vasopressin (AVP)-expressing neurons that contribute to synchronization within the SCN network (Mieda et al., 2015; Yamaguchi et al., 2013). Superimposed on this peptidergic division, nearly all SCN neurons are GABAergic, and one notable distinction between the two subregions lies in their response to γ-aminobutyric acid (GABA), particularly through GABA_A_ receptors. Although GABA typically exerts inhibitory effects by hyperpolarizing neurons, certain SCN neurons exhibit high intracellular chloride concentrations due to a day-length-dependent increase in the relative expression of the neuronal chloride importer (NKCC1) over the exporter (KCC2), causing GABA to have a depolarizing effect instead. This mechanism allows the two regions to maintain distinct stable phase gaps that together encode seasonal and photoperiodic information (Myung et al., 2015; Rohr et al., 2019).

One powerful paradigm for probing SCN plasticity is the use of T-cycles, light-dark cycles whose period (*T*) deviates from 24 h (e.g., T22 and T26 correspond to 11:11 LD and 13:13 LD, respectively). Similar to photoperiods, T-cycles induce aftereffects in the subsequent free-running period. Behavioral period generally shifts in the direction of the preceding T-cycle, with shorter T-cycles producing shorter behavioral periods and longer T-cycles producing longer ones. A key discrepancy emerges in explanted SCN recordings, which frequently show the opposite ordering: longer SCN periods after short T-cycles and shorter SCN periods after long T-cycles (Aton et al., 2004; Azzi et al., 2017; Molyneux et al., 2008). This inversion of aftereffects between behavior and explanted SCN tissue has motivated several theoretical efforts to reconcile the discrepancy and understand the encoding of light history within the SCN network.

Reduced two-oscillator descriptions of the *D*-and *V*-SCN, in which asymmetric attractive and repulsive coupling sets the phase relationship between subregions, were introduced alongside experimental work on photoperiodic encoding and were used to simulate jet lag, T-cycle entrainment, constant darkness, and constant light; in those treatments the intrinsic period of the *D* oscillator was assigned a fixed parameter for each condition (Myung et al., 2015; Myung & Pauls, 2018). One framework developed specifically to address the behavioral-explant discrepancy was proposed by Azzi et al. (2017). Their model represents the *V* and *D* subregions as two coupled phase oscillators with distinct intrinsic periods and asymmetric synchronizing and desynchronizing interactions. The model reproduced the experimentally observed phase separation between SCN subregions, with short T-cycles delaying the *D* oscillator relative to the *V* oscillator, and long T-cycles producing the opposite effect. It did not, however, reproduce the free-running period measured from SCN explants, and it relied on condition-specific parameter tuning and phase response functions, limiting its general applicability. Foster et al. (2019) subsequently proposed a fundamentally different explanation for the discrepancy between behavioral and explanted SCN aftereffects. In their model, the SCN consists of a heterogeneous population of oscillators whose intrinsic periods are distributed around 24 h. Oscillators with intrinsic periods closest to the entraining T-cycle exert the strongest influence on the network, resulting in persistent changes in population period. To simulate explant recordings, the model selectively removes highly connected oscillators, representing excitotoxic cell loss during slice preparation. This reproduces the inversion between behavioral and explanted SCN aftereffects observed in several T-cycle studies (Aton et al., 2004; Azzi et al., 2017; Molyneux et al., 2008). Yet the selective-cell-loss hypothesis does not readily explain why behavioral and SCN aftereffects remain largely consistent under photoperiod manipulations, particularly in studies using *Bmal1-ELuc* reporters (Myung et al., 2012, 2015). More recently, Kitaguchi et al. (2025) revisited the two-oscillator framework using a simplified asymmetric attractive-repulsive coupling model. Their analytical treatment demonstrated that repulsive coupling broadens the entrainment range and that slow relaxation near a phase-locked state can generate persistent T-cycle aftereffects following release into constant darkness. Nevertheless, the model primarily explains aftereffects as transient relaxation toward equilibrium and does not explicitly incorporate adaptive changes in intrinsic periods or coupling strengths that have been proposed to underlie longer-term photoperiodic encoding (Myung et al., 2015; Myung & Pauls, 2018; Rohr et al., 2019). Moreover, the model has not been evaluated under photoperiodic conditions and assumes that the measured SCN oscillator remains the principal determinant of behavioral period.

An alternative possibility is that behavioral timing *in vivo* emerges from interactions among multiple circadian oscillators distributed throughout the brain and peripheral tissues, including the choroid plexus, pineal gland, pituitary gland and peripheral organs (Myung et al., 2025a). Supporting this idea, transplantation experiments showed that replacing the SCN of animals exhibiting T-cycle aftereffects with an SCN from non-entrained donors does not abolish the behavioral aftereffect (Matsumoto et al., 1996), suggesting that circadian memory may not reside exclusively within the SCN. Furthermore, under extreme T-cycle entrainment, animals can exhibit two behavioral rhythmic components: one synchronized to the imposed light cycle and another free-running component whose period varies inversely with T-cycle length (Campuzano et al., 1998). One interpretation is that the explanted SCN represents the oscillator exhibiting the opposite period shift, whereas another, as yet unidentified, circadian oscillator or downstream output pathway remains positively coupled to the entraining cycle and ultimately determines behavioral timing. Alternatively, the intrinsic period expressed by the SCN *in vivo* may itself be shaped by reciprocal interactions with extra-SCN oscillators, such that isolation *ex vivo* reveals a different dynamical state. These possibilities challenge the common assumption that the explanted SCN necessarily represents the behaviorally relevant circadian pacemaker.

We therefore hypothesized that the behavioral period *in vivo* is generated through interactions between the SCN and an additional circadian oscillator, denoted *X*. *X* is not assigned to a specific anatomical structure but represents the combined influence of unexplanted SCN components, extra-SCN clocks, peripheral oscillators, or systemic circadian signals that can modify SCN dynamics *in vivo*. Under this framework, the SCN and *X* jointly determine behavioral timing while they remain coupled in the intact animal. Explantation removes systemic feedback from *X*, allowing the isolated SCN to relax toward a different dynamical state. The behavioral and *ex vivo* SCN periods may therefore differ after the same light history without requiring either measurement to be considered erroneous. Reporter-specific sampling of dorsal, ventral, or molecularly distinct SCN populations may further influence the period observed *ex vivo*.

To examine this hypothesis, we first compiled matched measurements of behavioral and explanted SCN free-running periods following photoperiod and T-cycle entrainment from the published literature. We then developed a phase model comprising *D* and *V* SCN oscillators coupled to a putative systemic oscillator *X*. Light-history memory adaptively modified the intrinsic periods of the *D* SCN oscillator and *X*. In contrast to earlier two-oscillator treatments, here it evolves as a dynamic state variable driven by accumulated light history. The complete coupled system was used to simulate behavioral period *in vivo*, whereas *ex vivo* SCN dynamics were simulated by removing feedback from *X* at the time of release into constant darkness. The model was fitted to behavioral periods and *D*-*V* phase relationships across photoperiod and T-cycle conditions. We further compared intrinsic-period adaptation with an alternative coupling-adaptation mechanism and generated predictions for previously untested lighting schedules.

## Methods

### Literature extraction and data curation

To establish quantitative targets for model development, we first compiled published measurements of behavioral free-running period and SCN period following non-standard light exposure. The initial literature set was assembled from the reference lists of Azzi et al. (2017) and Myung et al. (2015), and was supplemented by Google Scholar searches for studies reporting behavioral and SCN rhythms under photoperiod or T-cycle manipulations. Studies were included when behavioral period, SCN period, or both could be extracted together with sufficient information to classify the lighting paradigm as standard photoperiod, T-cycle, or photoperiod × T-cycle interaction. Period estimates were obtained directly from texts, tables, or digitized from figures using WebPlotDigitizer v5 (Automeris; https://automeris.io/). All extracted measurements were harmonized into a common long-format database containing study, lighting paradigm, reporter, age group, SCN region, measurement type, condition, mean period, SEM, and sample size where available. To avoid over-representing studies reporting multiple SCN regions from the same experimental cohort, one representative SCN measurement was selected for each study, reporter, age group, and condition using a predefined anatomical priority of whole SCN, dorsal SCN, and posterior SCN. Behavioral and SCN periods from matching experimental groups were then paired for subsequent comparison. These curated behavioral aftereffect measurements served as quantitative targets for computational model fitting.

## Computational Modeling

### Model overview

The SCN was represented by coupled dorsal (*D*) and ventral (*V*) phase oscillators. A third oscillator, *X*, represented the net influence of extra-SCN or systemic circadian signals that remain coupled to the SCN *in vivo*. The same equations and parameter set were used for all lighting schedules. *In vivo* behavior was simulated with the complete *D*-*V*-*X* network, whereas *ex vivo* SCN dynamics were simulated from the same entrained state after feedback from *X* to the SCN was removed. This design tested whether loss of systemic feedback at explantation could reveal an SCN aftereffect different from the period governing behavior *in vivo*.

### Light schedules and simulation protocol

Each simulation comprised 30 days under the specified entrainment schedule followed by 14 days of constant darkness (DD). Short photoperiod 8:16 LD (SP), standard light-dark 12:12 LD (LD), and long photoperiod 16:8 LD (LP) schedules had 24 h cycles containing 8, 12, and 16 h of light, respectively. T22 and T26 contained 11 and 13 h of light per cycle. The binary light state was denoted *I*(*t*), with *I*(*t*) = 1 during light and *I(t*) = 0 during darkness. The phase of the external schedule was:

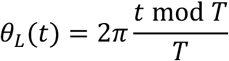

where *T* is the duration of the entraining cycle. The differential equations were integrated by the forward Euler method with a 0.25-h time step.

### Oscillator phases and behavioral output

The composite SCN phase was defined as the weighted circular mean of the dorsal and ventral oscillators,

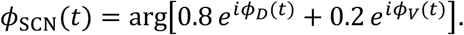

Oscillator activity was represented as *y_j_*(*t*) = −sin[*φ_j_*(*t*)], and behavioral activity was defined as *y*_beh_(*t*) = −sin[*φ*_SCN_(*t*)]. Because mice are nocturnal, activity was represented such that its peak occurred during the dark phase. The 80:20 weighting assumes that behavioral timing is primarily determined by the dorsal SCN while retaining a smaller contribution from the ventral light-responsive oscillator (Keene et al., 2026). The systemic oscillator *X* influenced behavior only indirectly through its coupling to the SCN. The *D*-*V* phase difference was defined as *ϕ_DV_*(*t*) = wrap(*ϕ_D_*(*t*) − *ϕ_V_*(*t*)), with wrapping to (−*π*, *π*]. Positive values indicate that the dorsal oscillator leads the ventral oscillator. Phase differences were converted to hours as

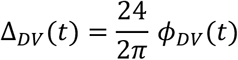

### Light-history memory and adaptive intrinsic periods

Separate light-history memories were assigned to *D* and *X*. For *j* ∈ {D, X}, subjective day was represented by

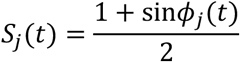

and memory evolved according to

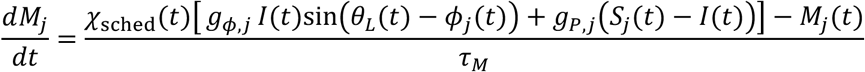

Here *χ*_sched_(*t*) = 1 during scheduled entrainment and *χ*_sched_(*t*) = 0 in DD. Both memories used *τ*_M_ = 240 h. Consequently, after release into DD, *dM_j_*/*dt* = −*M_j_*/*τ*_M,_ corresponding to a 10-day 1/*e* decay time. The phase-sensitive term encodes the circadian phase at which light is received, whereas the photoperiod term encodes disagreement between the external light state and subjective day.

The memory states modified the intrinsic periods of *D* and *X*:

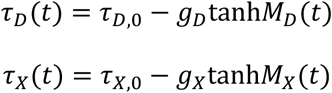

Adaptive periods were bounded numerically between 16 and 34 h, whereas *τ_V_* remained fixed. Photoperiod agreement between *X* and the SCN was imposed at the output level: *X*, *D*, and *V* were required to retain the *in vivo* ordering LP < LD < SP and comparable SP-LD and LD-LP contrasts. Identical photoperiod-memory coefficients were not required because the memory terms for *D* and *X* are evaluated at different oscillator phases.

### Coupled phase-oscillator equations

The following describes the coupled dynamics of the *D* (*ϕ*_D_), *V* (*ϕ*_V_) and *X* (*ϕ*_X_) oscillators driven by light (*I*) and global SCN output (*ϕ*_SCN_).

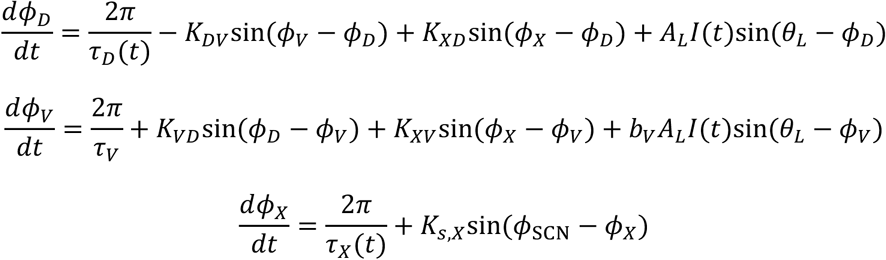

Under this sign convention, *K_VD_* is the attractive *D*-to-*V* term and *K_DV_* contributes a repulsive *V*-to-*D* interaction. Light acts directly on both SCN oscillators, with the ventral light response scaled by *b_V_*. *X* receives no direct light input. Its adaptive memory is nevertheless driven by a phenomenological representation of light history, corresponding to indirect systemic access to environmental timing information (through interaction with the SCN); its schedule dependence arises through *M*_X_ and coupling from the composite SCN phase.

### In vivo and ex vivo simulations

The complete *D*-*V*-*X* system was simulated throughout entrainment. For the *in vivo* condition, all interactions were retained after release into DD. For the *ex vivo* condition, the phases and memory states reached at DD onset were preserved, but feedback from *X* to the SCN was removed by setting *K_XD_* = *K_XV_* = 0. The isolated *D*-*V* subsystem therefore carried the state acquired *in vivo* but subsequently evolved without systemic feedback. *Ex vivo* summaries were restricted to *D* period, *V* period, and the *D*-*V* phase gap.

### Period and phase-gap estimation

Periods were estimated from the complete 14-day DD record (DD hours 0–336); no initial DD days were discarded. Lomb-Scargle periodograms were computed with the Astropy implementation over an 18–30 h search range, using 20 frequency samples per expected periodogram peak. The reported period was the inverse of the frequency with maximum power. The quantitative *D*-*V* phase gap was the arithmetic mean of the wrapped gap over DD hours 0–168, corresponding to the first seven DD days. This window matched the early-DD interval used in the source phase-gap datasets.

### Fitting targets and parameter estimation

The *in vivo* behavioral targets are summarized in **Table 2**. Behavioral period was used as the primary fitting target because behavioral aftereffects were more consistent across studies, whereas reported *ex vivo* SCN periods varied with the molecular reporter (**Figure 1**). *Ex vivo* SCN periods and *D*-*V* phase gaps were therefore used as secondary constraints on the explant SCN dynamics. Because the SP observations were compatible with near-synchrony, no strict positive lower bound was imposed on the SP phase gap during final model selection.

**Figure 1.**
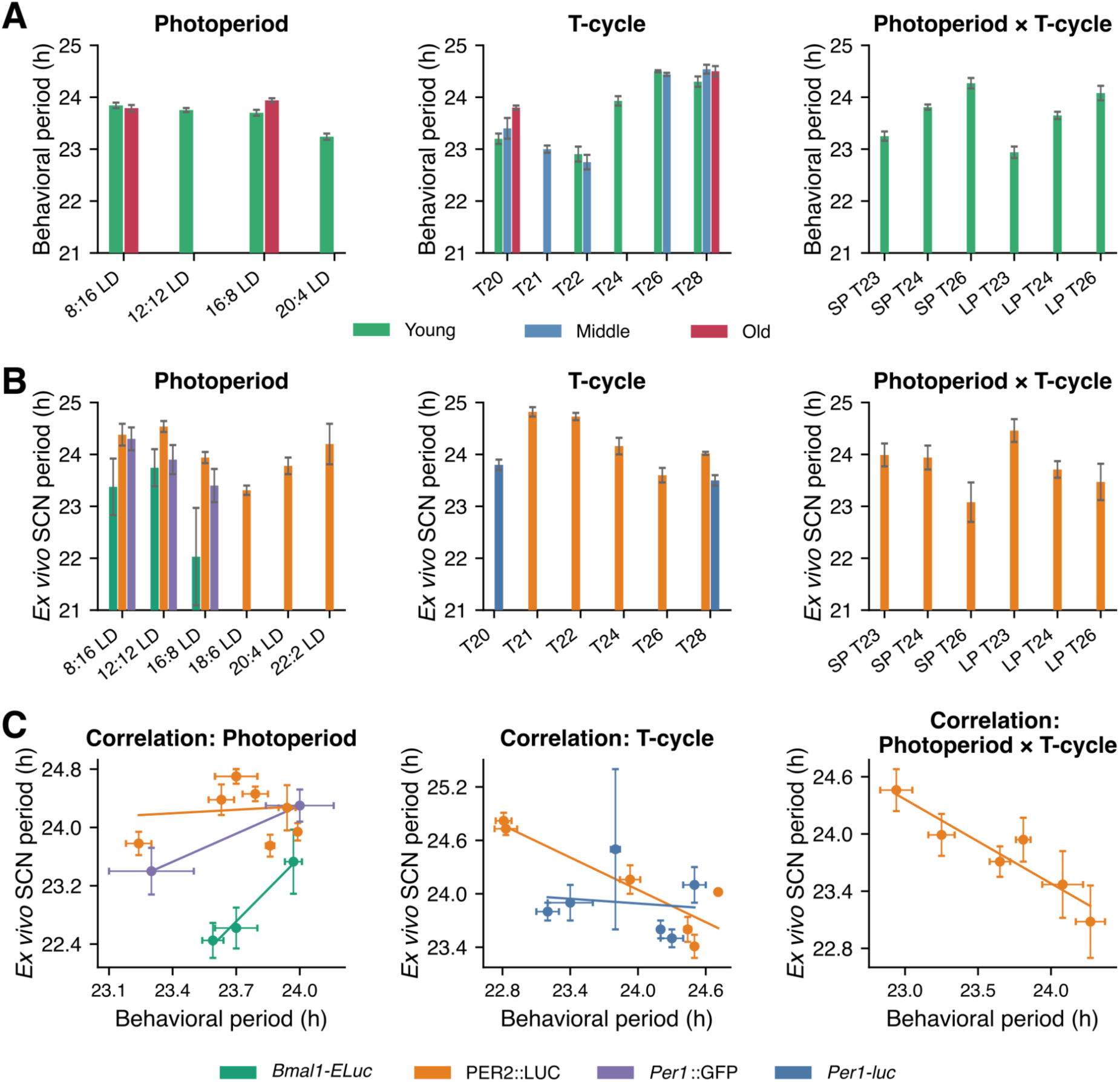
Relationship between behavioral and SCN circadian period across photoperiod, T-cycle, and combined entrainment paradigms. **(A)** Mean behavioral free-running period measured following entrainment to different photoperiods (left), T-cycles (middle), and combined photoperiod × T-cycle schedules (right). Data are shown for the available age groups (young, middle-aged, and old, where applicable). **(B)** Corresponding SCN circadian periods measured from *ex vivo* slice recordings under the same entrainment paradigms using different molecular reporters. Photoperiod data include *Bmal1-ELuc*, PER2::LUC, and *Per1*::GFP reporters, whereas T-cycle and photoperiod × T-cycle datasets are primarily derived from PER2::LUC and *Per1-luc* recordings. Bars represent mean ± SEM. **(C)** Correlation between behavioral and SCN circadian periods across studies. Left, photoperiod datasets separated by reporter; center, T-cycle datasets; right, combined photoperiod × T-cycle datasets. Each point represents an experimental condition, with horizontal and vertical error bars indicating SEM for behavioral and SCN period, respectively. Linear regression lines are shown for each reporter. Behavioral and SCN periods exhibited reporter-dependent relationships across entrainment paradigms, with PER2::LUC showing a positive association under photoperiod entrainment but a negative association under T-cycle and combined photoperiod × T-cycle paradigms, whereas *Bmal1-ELuc* displayed a positive relationship with behavioral period. This divergence highlights that distinct molecular oscillatory readouts capture different aspects of SCN network adaptation to environmental light schedules.

For SP, LP, and LD, *ex vivo* SCN periods were constrained to remain near their corresponding *in vivo* SCN periods. The objective also included the *in vivo* photoperiod contrasts SP-LD = 0.27 h and LD-LP = 0.11 h. For T-cycle conditions, the *ex vivo* period targets followed the inverse aftereffect pattern: 24.50 h after T22 and 22.83 h after T26. Parameter estimation proceeded in two stages. A seeded Latin-hypercube search broadly sampled the parameter space, after which the leading candidates were refined by finite-difference local gradient descent. Candidate simulations were evaluated in parallel with Numba. During screening, period was approximated from the 14-day slope of unwrapped phase to reduce computational cost. The leading candidates and one mechanistically calibrated architecture candidate were then re-simulated, and their periods were recomputed with the exact 14-day Lomb-Scargle periodogram. Final selection used a diagnostic-compromise score combining behavioral-period, *ex-vivo*-period, and first-seven-day phase-gap residuals with penalties enforcing the prespecified period orderings and phase-gap signs. The raw screening objective and the final Lomb-Scargle score were treated as distinct quantities. The search used seed 1729, 65,536 Latin-hypercube samples, 32 local starts, 55 local iterations per start, and a batch size of 1,024. Selected model parameters and optimization scores were shown in **Table S1**.

### Reduced D-V phase-locking and bifurcation analysis

After removal of light input and systemic feedback, the *D*-*V* phase difference was approximated by

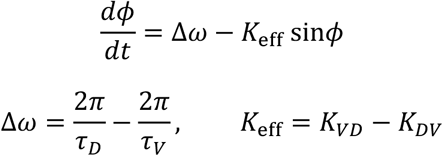

A fixed point exists when |*Δω/K*_eff_| ≤ 1 (Myung & Pauls, 2018). The reduced locking ratio is defined as *R* = |*Δω/K*_eff_|; *R* > 1 indicates that no phase-locked fixed point exists in the reduced system. Stability additionally requires *K*_eff_ cos(*φ*\*) > 0. Stable and unstable equilibria coalesce at |*Δω/K*_eff_| = 1 via a saddle-node bifurcation (Myung et al., 2025b). Bifurcation curves were generated by varying *K_VD_* while holding *K_DV_* and the condition-specific mean early-DD value of *τ_D_* fixed. Solid and dashed branches denote stable and unstable equilibria, respectively, and shaded regions indicate the absence of a reduced fixed point (**Figure 5A**).

### Comparison of intrinsic-period and coupling adaptation

To test whether light-history-dependent responses were more naturally encoded through intrinsic period or *D*-*V* coupling, two mutually exclusive architectures were simulated. In the period-adaptive architecture, memory modified *τ_D_* while *K*_eff_ remained fixed:

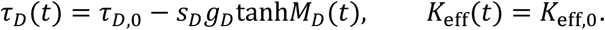

In the coupling-adaptive architecture, *τ_D_* and *τ_V_* were fixed and the same memory signal modified *K*_eff_:

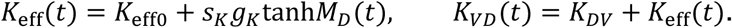

The memory trajectory *M*_D_(*t*), light schedules, initial phases, and all non-adaptive parameters were identical between architectures. The coupling gain *g*_K_ was estimated by a one-dimensional grid search against the five *ex vivo* phase-gap targets weighted by their SEM. The coupling-adaptive model was therefore treated as a mechanistic comparator rather than as a fully re-optimized competing model.

The dimensionless gain scales *s*_D_ and *s*_K_ were varied independently from 0 to 2; *s* = 0 removed the corresponding adaptation and *s* = 1 reproduced its selected reference gain. Full-model central differences used *s* = 0.95 and 1.05 and were normalized by the resulting schedule-specific change in *Δω* or *K*_eff_. *D* and *V* periods were estimated from the complete 14-day DD record, phase gaps from the first-seven-day DD mean, and locking from *R* = |*Δω*/*K*_eff_|. Analytical derivatives of the reduced locked model are given in the Supplementary Information.

After fitting, the selected model was also simulated without further parameter adjustment under 18:6 LD, T23, and T25 using the same entrainment, DD, and *in vivo*/*ex vivo* release procedures.

## Results

### Descriptive synthesis of behavioral and SCN period relationships

We compiled 30 matched behavioral-SCN period pairs comprising 12 photoperiod, 12 T-cycle, and 6 photoperiod × T-cycle conditions. Across the extracted literature, behavioral aftereffects generally followed the expected direction of the entraining light schedule, whereas SCN period aftereffects varied considerably depending on the molecular reporter and entrainment paradigm (**Figure 1A, B**). In particular, photoperiod studies showed relatively consistent behavioral period changes but heterogeneous SCN responses across reporters, while T-cycle studies revealed an apparent dissociation between behavioral and SCN period aftereffects.

To quantify these relationships, behavioral and SCN periods were analyzed separately for each reporter within each condition family (**Table 1**; **Figure 1C**). Under photoperiod entrainment, the largest dataset (PER2::LUC, *n* = 8) showed no clear association between behavioral and SCN periods (*r* = 0.10, *p* = 0.82). *Bmal1-ELuc* and *Per1*::GFP exhibited positive trends, although these were based on three and two matched condition means, respectively. Reporter differences are also apparent in absolute periods. *Bmal1-ELuc* explant period did not differ significantly from behavioral free-running period in either anterior or mid-to-posterior SCN, whereas PER2::LUC explant period was significantly longer in both (Myung et al., 2012)

**Table 1.** Reporter-specific correlations between behavioral and SCN free-running periods across photoperiod and T-cycle entrainment paradigms.

| Condition family | Reporter | *n* pairs | Pearson *r* | *p* value | Slope, SCN vs behavior | Reference |
| --- | --- | --- | --- | --- | --- | --- |
| Photoperiod | *Bmal1-ELuc* | 3 | 0.99 | 0.088 | 2.94 | (Myung et al., 2015) |
| Photoperiod | PER2::LUC | 8 | 0.097 | 0.8181 | 0.16 | (Buijink et al., 2020; Evans et al., 2013; Myung et al., 2012) |
| Photoperiod | *Per1::GFP* | 2 | 1.00 | not tested | 1.29 | (Ciarleglio et al., 2011) |
| T-cycle | *Per1-luc* | 6 | −0.13 | 0.8110 | −0.089 | (Aton et al., 2004) |
| T-cycle | PER2::LUC | 6 | −0.90 | 0.0147 | −0.61 | (Azzi et al., 2017;<br>Molyneux et al.,<br>2008) |
| Photoperiod<br>× T-cycle | PER2::LUC | 6 | −0.9325 | 0.0067 | −0.88 | (Tackenberg et<br>al., 2020) |

**Notes.** Pearson correlations were calculated using matched behavioral and SCN free-running period measurements from the same study, reporter, age group, and entrainment condition. The regression slope represents the change in SCN period (h) per 1-h change in behavioral period. *p*-values were not calculated for *n* = 2 because Pearson correlation has no residual degrees of freedom.

**Table 2.**
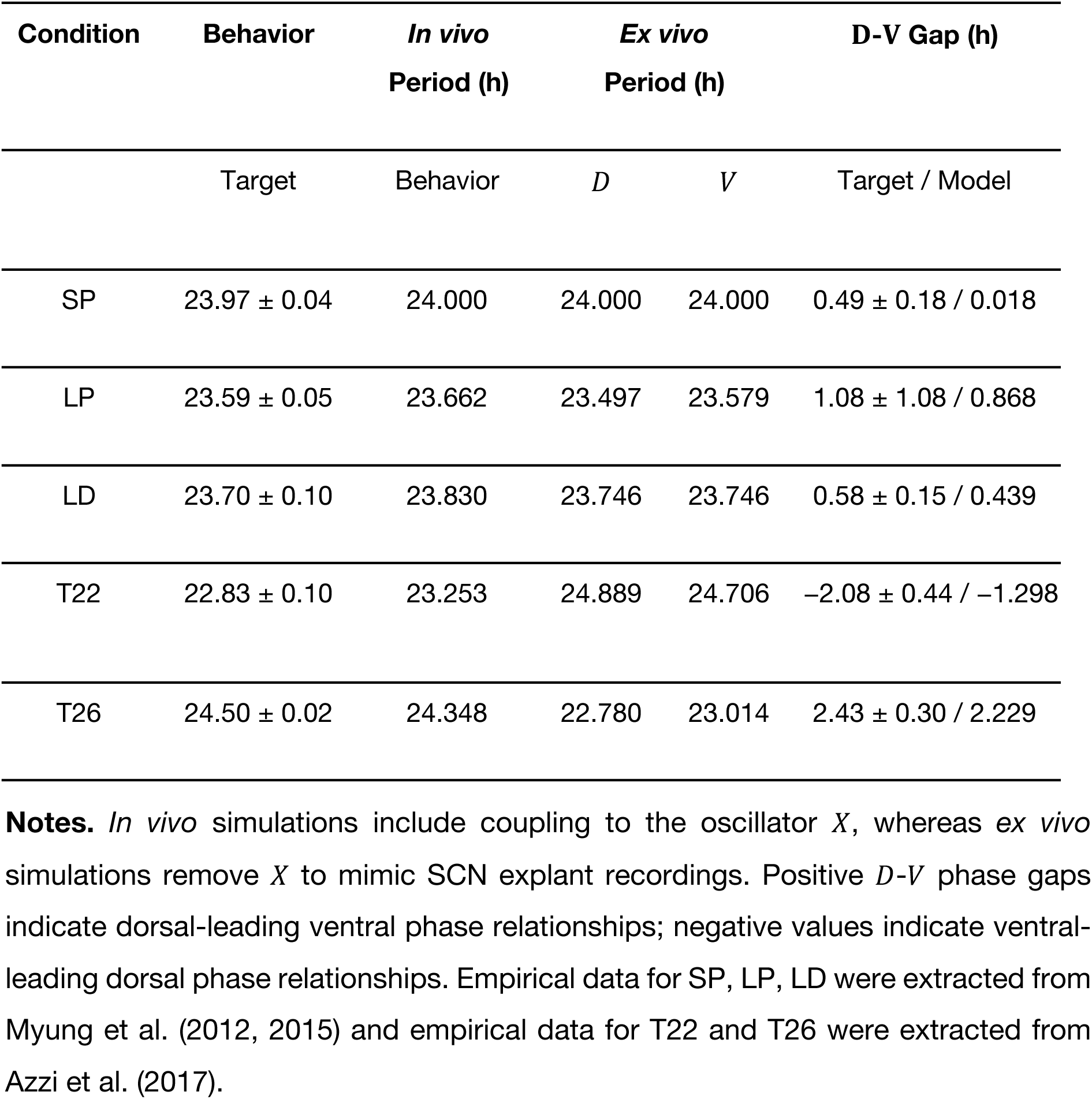
Fitted behavioral, oscillator-period, and D-V phase-gap outputs across light schedules.

In contrast, T-cycle datasets displayed a reporter-dependent pattern. PER2::LUC measurements showed a strong negative correlation between behavioral and SCN periods (*r* = −0.90, \**p* = 0.015), indicating that conditions producing longer behavioral periods were associated with shorter SCN periods. This relationship was weaker in the *Per1-luc* dataset (*r* = −0.13, *p* = 0.81). Similarly, the combined photoperiod × T-cycle dataset exhibited a strong negative correlation in PER2::LUC (*r* = −0.93, \*\**p* = 0.0067).

These findings indicate that the relationship between behavioral and *ex vivo* SCN aftereffects depends on both the entrainment schedule and the molecular reporter. They therefore challenge the assumption that the free-running period measured from an SCN explant necessarily reflects the circadian dynamics governing behavior in the intact animal. To examine whether this discrepancy could arise from the loss of systemic feedback after explantation, we next developed a *D*-*V* SCN model coupled to an additional oscillator, *X*. The complete *D*-*V*-*X* system was used to simulate behavioral timing *in vivo*, whereas *ex vivo* SCN dynamics were simulated by removing feedback from *X* at the onset of DD. We then tested whether a single parameter set could reproduce the behavioral periods, *ex vivo* SCN periods, and *D*-*V* phase relationships across photoperiod and T-cycle conditions.

### Systemic feedback reconciled *in vivo* behavior with *ex vivo* SCN aftereffects

The selected *D*-*V*-*X* model reproduced the expected *in vivo* ordering under both photoperiod and T-cycle entrainment (**Figure 2**; **Table 2**). Behavioral periods followed LP < LD < SP (23.662, 23.830, and 24.000 h) and T22 < LD < T26 (23.253, 23.830, and 24.348 h). Because *D*, *V*, and *X* remained strongly coupled *in vivo*, their estimated periods closely matched the composite behavioral period within each schedule.

**Figure 2.**
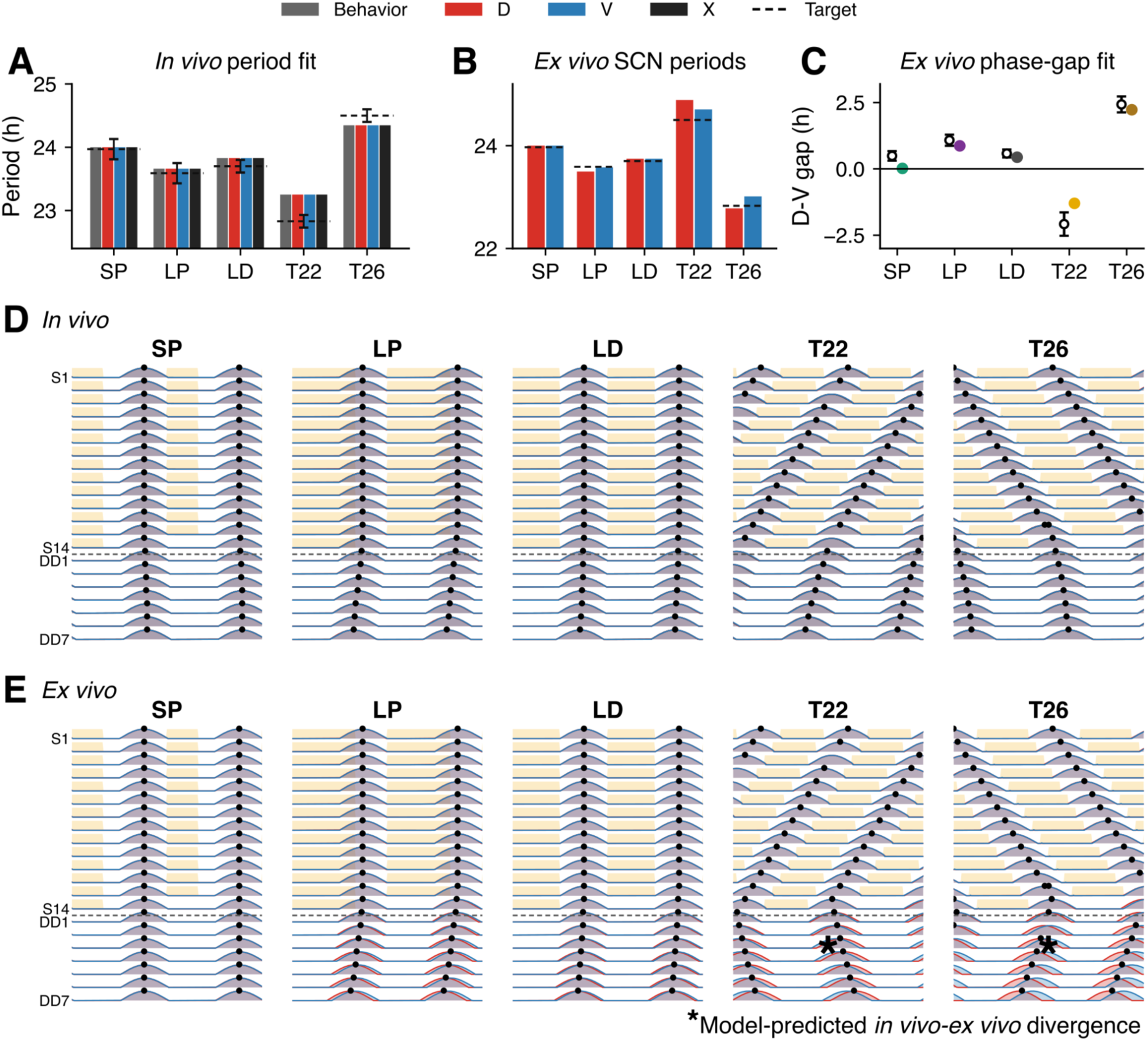
Model fits and simulated oscillator dynamics across lighting schedules. **(A)** Simulated *in vivo* free-running periods following entrainment under short photoperiod (SP), long photoperiod (LP), standard 12:12 light-dark (LD), T22, and T26 schedules. Bars show the behavioral rhythm (gray), dorsal SCN oscillator (*D*, red), ventral SCN oscillator (*V*, blue), and systemic oscillator (*X*, black). Dashed lines indicate the behavioral target periods used for model fitting. **(B)** Predicted *ex vivo* free-running periods of the dorsal and ventral SCN oscillators after removal of oscillator *X* input. Dashed lines indicate the expected SCN tissue period under each condition. **(C)** Comparison between empirical (open circles) and simulated (filled circles) *D*-*V* phase differences measured on the first day in constant darkness after explant preparation. Error bars represent the experimental uncertainty of the empirical measurements. **(D–E)** Double-plotted oscillator activity during the final 14 days of entrainment (S1–S14) followed by the first 7 days in constant darkness (DD1– DD7). Yellow shading denotes the scheduled light phase before release into constant darkness. **(D)** *In vivo* simulations showing behavioral peaks (black dots) derived from the composite SCN output. **(E)** Corresponding *ex vivo* simulations after disconnection of the systemic oscillator, illustrating the intrinsic dynamics of the *D* (red) and *V* (blue) SCN oscillators during the aftereffect period.

Removing feedback from *X* at DD onset exposed a different SCN response after T-cycle entrainment. *Ex vivo D* and *V* periods were longer after T22 (24.889 and 24.706 h) but shorter after T26 (22.780 and 23.014 h), thereby reversing the *in vivo* T-cycle ordering. Photoperiod aftereffects remained in the same direction *in vivo* and *ex vivo*. The simulated *D*-*V* phase gap was positive after SP, LP, LD, and T26, indicating that *D* led *V*, but was negative after T22, indicating that *V* led *D*. The largest remaining gap residual occurred after T22, for which the simulated value was −1.298 h compared with the target of −2.08 h. SP approached synchrony, with a simulated gap of 0.018 h.

### Reduced phase dynamics distinguish stable photoperiod phase relationships from evolving phase dynamics in T-cycle aftereffects

The local weighted-SSE surface placed the selected parameter set within a low-error region of the dorsal phase-and photoperiod-memory gain landscape (**Figure 3A**). This plotted SSE was used as a transparent residual summary and was not identical to the constrained fitting objective used during model selection.

**Figure 3.**
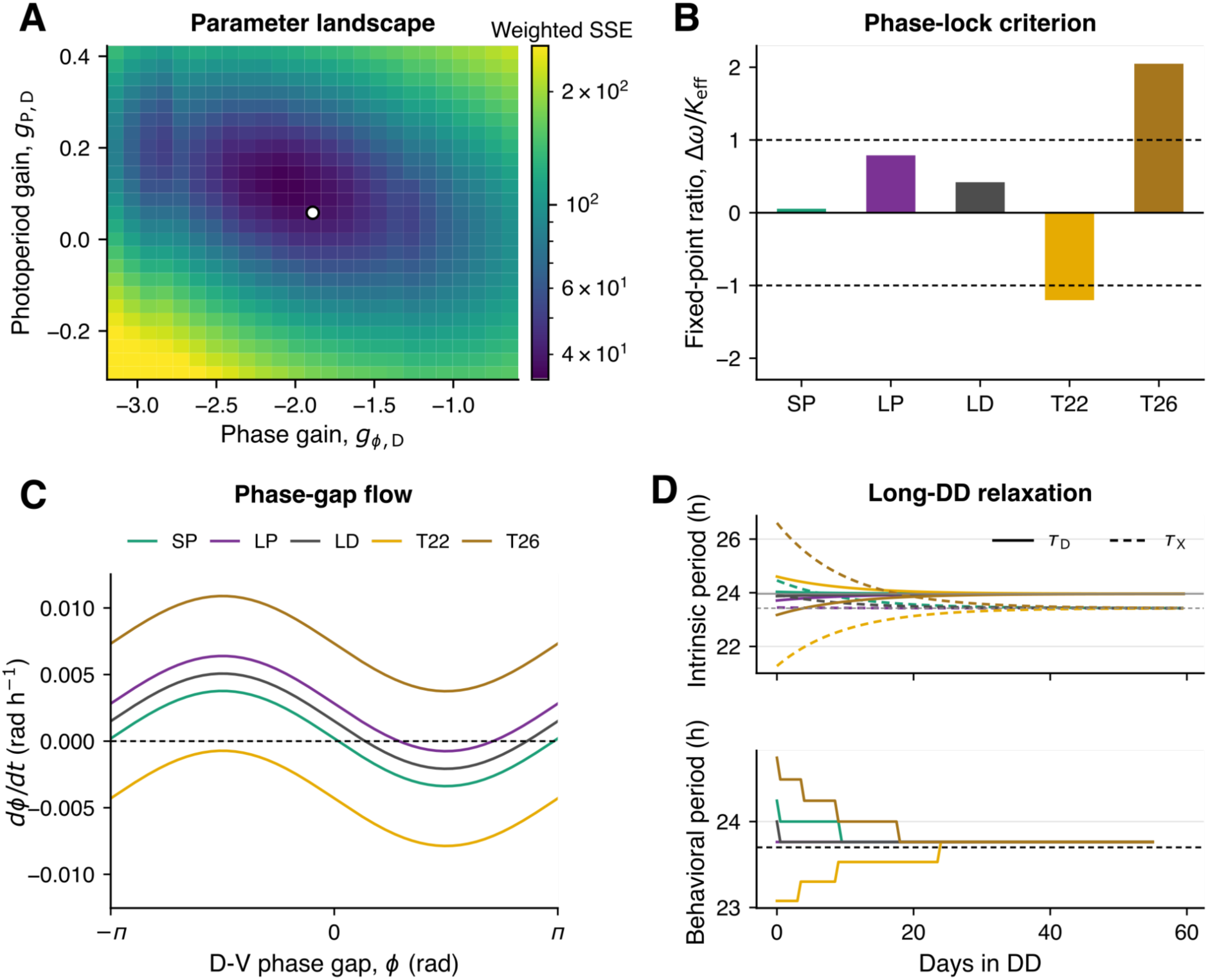
Parameter landscape, phase-locking analysis, and relaxation dynamics. **(A)** Local parameter landscape showing the weighted sum-of-squared errors (SSE) as a function of the phase memory gain (*g_ϕ,D_*) and photoperiod memory gain (*g_P,D_*). The white circle indicates the selected parameter set. The displayed SSE summarizes the fit to experimental observations and is intended for visualization; it is not identical to the constrained optimization objective used during parameter fitting. **(B)** Reduced fixed-point ratio, *Δω*/*K*_eff_, for each lighting condition. Values satisfying |*Δω*/*K*_eff_| ≤ 1 (dashed lines) permit stable fixed points in the reduced time-independent system; their stability additionally depends on *K*_eff_ cos*ϕ^∗^* > 0. **(C)** Phase-gap flow curves showing *dϕ/dt* as a function of the *D*-*V* phase difference (*ϕ*) under each lighting condition. Intersections with the zero line correspond to phase-locked steady states, with the local slope determining their stability. **(D)** Relaxation of the intrinsic periods of the dorsal SCN oscillator (*τ_D_*, solid lines) and systemic oscillator (*τ_X_*, dashed lines) following release into DD (top), together with the resulting behavioral free-running period (bottom).

At the fitted operating point, *K*_eff_ was 0.003575 rad h⁻¹ for all schedules, whereas *Δω* depended on the schedule-specific early-DD value of *τ_D_* (**Table S2**). The reduced criterion predicted phase locking for SP (*R* = 0.055), LP (*R* = 0.789), and LD (*R* = 0.420), but not for T22 (*R* = 1.202) or T26 (*R* = 2.049) (**Figure 3B**). The phase-flow curves showed corresponding zero crossings for the photoperiod conditions, whereas the T-cycle conditions lay outside the instantaneous reduced locking range (**Figure 3C**). This reduced analysis is an early-DD diagnostic rather than a replacement for the full time-dependent simulation. Both *M_D_* and *M_X_* continued to decay in DD, causing *τ_D_*, *τ_X_*, *Δω*, and the locking margin to evolve. With *τ*_M_ = 240 h, the memory states relaxed over a 10-day time scale. Without any condition-specific release reset, long-DD simulations converged toward an approximately 23.76 h behavioral period. LD and SP approached this asymptote by approximately DD day 1, whereas the other light schedules relaxed more gradually (**Figure 3D**).

### Reduced model analysis and full model simulations support intrinsic period adaptation as a parsimonious control mechanism for light adaptation

Reduced phase analysis was used to interpret two mutually exclusive mechanisms by which light-history memory could modify SCN dynamics: adaptation of the dorsal intrinsic period or adaptation of *D*-*V* coupling (**Figure 4; Tables S3 and S4**). The analytical derivation is provided in the Supplementary Information and was used to mechanistically interpret the results of the full model simulations.

**Figure 4.**
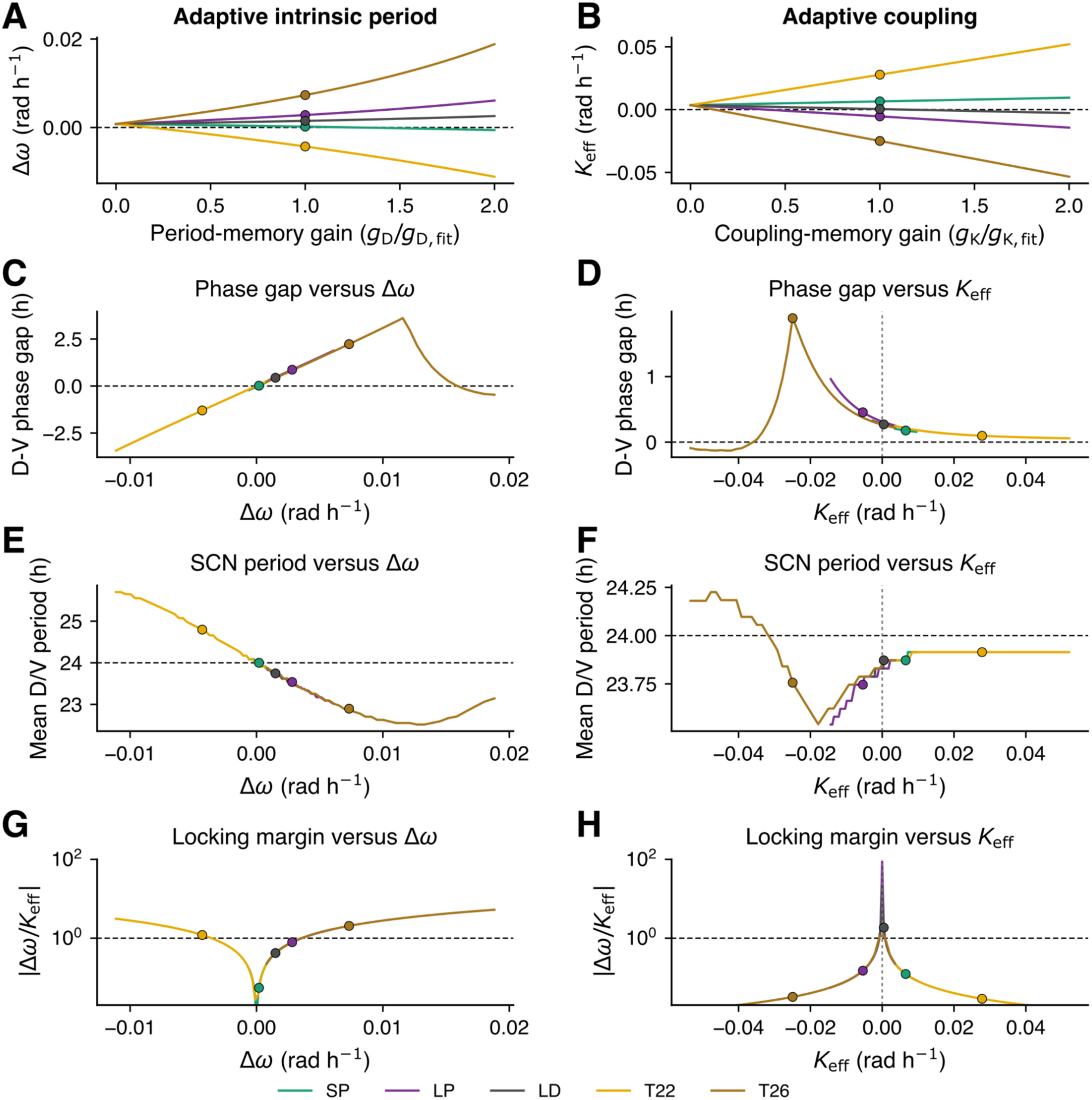
Mutually exclusive intrinsic-period and coupling-adaptation mechanisms. **(A)** Condition-specific frequency mismatch *Δω* generated when light-history memory adapts *τ_D_* and *K*_eff_ remains fixed. **(B)** Condition-specific *K*_eff_ generated when the same memory adapts coupling while both intrinsic periods remain fixed. Horizontal axes show each adaptive gain scaled from zero to twice its selected value; black-edged points mark *s*_D_ = *s*_K_ = 1. **(C, D)** Mean *D*-*V* phase gap during the first seven DD days plotted against the direct driver altered by each architecture. **(E, F)** *D*-and *V*-SCN periods estimated from the complete 14-day DD record. **(G, H)** Reduced locking ratio *r* = |*Δω*/*K*_eff_|. The horizontal dashed line marks *r* = 1, above which the reduced model has no phase-locked fixed point. The vertical reference in the coupling panels marks *K*_eff_ = 0.

**Figure 5.**
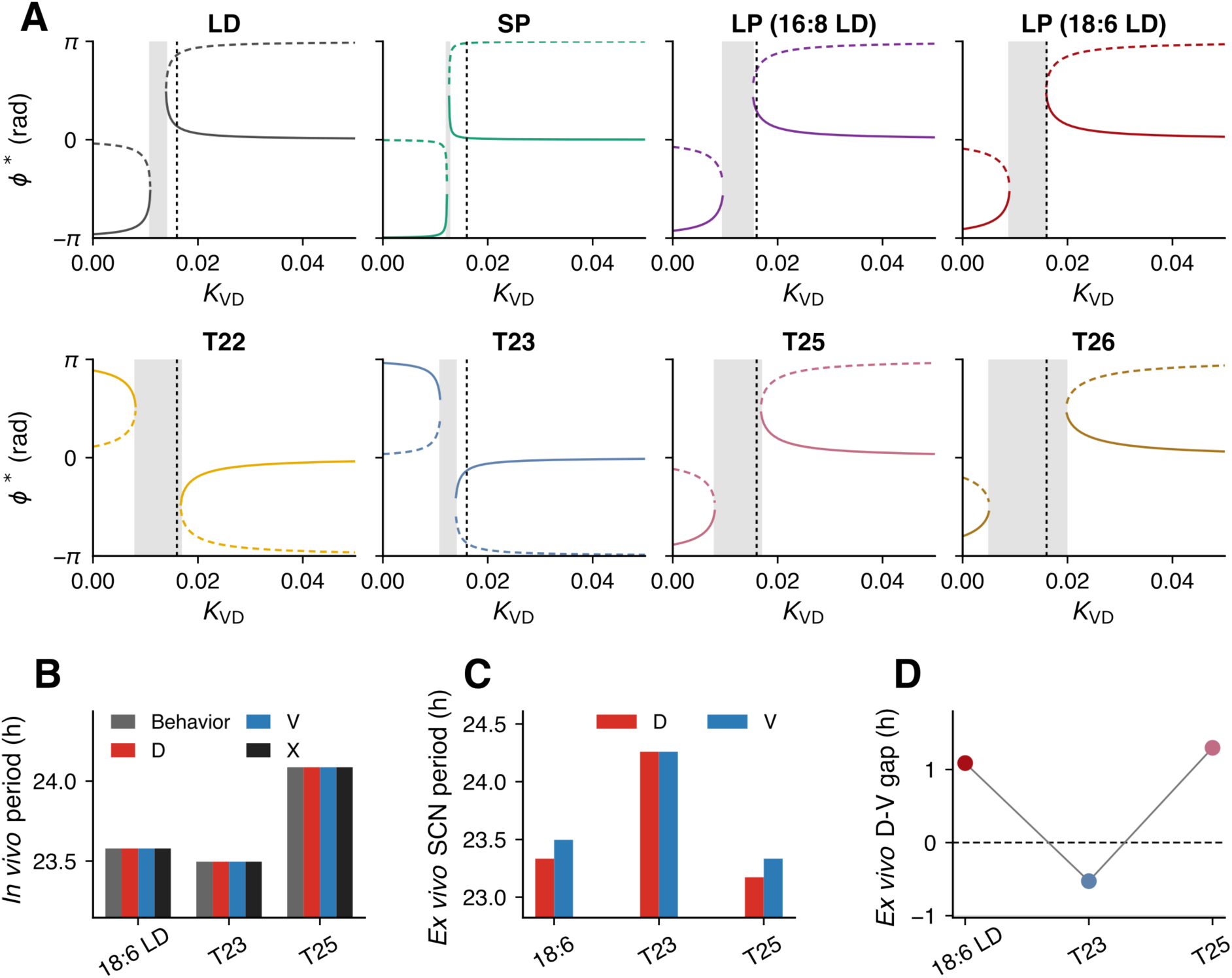
Bifurcation analysis and model predictions for untested lighting schedules. **(A)** Equilibrium phase differences (*ϕ^∗^*) as a function of coupling terms *K_VD_* with *K_DV_* held constant. Panels show the fitted conditions (LD, SP, LP, T22, and T26) together with model predictions for 18:6 LD, T23, and T25. The gray shaded region corresponds to parameter values for which the phase-locking criterion |*Δω/K*_eff_| > 1 is violated and no phase-locked equilibrium exists. |*Δω/K*_eff_| = 1 marks saddle node bifurcation boundaries. Solid and dashed branches denote stable and unstable equilibria, respectively. Vertical dashed lines indicate the fitted *K_VD_* value used in the simulations. **(B)** Predicted *in vivo* free-running periods of the behavioral rhythm (gray), dorsal SCN oscillator (*D*, red), ventral SCN oscillator (*V*, blue), and systemic oscillator (*X*, black) following entrainment to 18:6 LD, T23, and T25 schedules. **(C)** Predicted *ex vivo* free-running periods of the dorsal and ventral SCN oscillators after removal of systemic input. **(D)** Predicted *D*-*V* phase differences measured on the first 7 days of *ex vivo* culture following entrainment under the novel light schedules.

In the adaptive period architecture, light-history memory altered the sign and magnitude of the intrinsic frequency mismatch, Δ*ω*, while *K*_eff_ remained fixed (**Figure 4A**). In the full simulations, SP, LP, LD, T22, and T26 occupied different positions along approximately shared phase-gap, SCN-period, and locking-ratio trajectories as *Δω* changed (**Figure 4C,E,G**). Within the stationary range of the reduced model, |Δ*ω/K*_eff_| < 1, the equilibrium phase gap varies smoothly with Δ*ω,* although its sensitivity increases near the fixed-point boundary. Intrinsic-period adaptation therefore provided a common frequency-mismatch axis through which different light histories could be represented continuously without changing the underlying *D*-*V* coupling.

In the adaptive coupling architecture, Δ*ω* remained fixed and light-history memory instead generated schedule-dependent changes in *K*_eff_ (**Figure 4B**). The response depended on the reduced ratio: 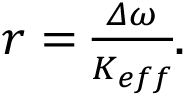 Coupling changes had comparatively modest effects when |*r*| was small but became strongly nonlinear as |*r*| approached 1. Varying *K*_eff_ could also move the system into a region with no stationary *D*-*V* phase relationship or through *K*_eff_ = 0, where the effective interaction vanished and changed sign. Coupling adaptation therefore modified not only the magnitude of the phase difference but also the existence and branch structure of stationary *D*-*V* phase organization.

The common-period derivatives provided a similar interpretation. Where a stable *D*-*V* state existed, sensitivity to *Δω* was direct, producing an approximately shared period relationship across schedules (**Figure 4E**). By contrast, the effect of changing *K*_eff_ was proportional to the frequency mismatch. When |Δ*ω/K*_eff_| was small, changes in coupling produced little change in the stationary period, explaining why the curves in **Figure 4F** were comparatively flat over much of their range, whereas changing *Δω* produces a clearer and more continuous period response in **Figure 4E**.

The phase-gap-weighted grid search selected *g*_K_ = −0.484 rad h⁻¹ (weighted SSE = 43.799). Nevertheless, the coupling-only architecture failed to reproduce the negative T22 phase gap: the simulated T22 gap was +0.097 h rather than the targeted −2.08 h. The other simulated gaps were 0.177 h for SP, 0.455 h for LP, 0.271 h for LD, and 1.893 h for T26. This failure arose because the intrinsic frequency difference between the *D* and *V* oscillators remained fixed. Changing only the coupling strength could modify the magnitude of the *D*-*V* phase difference but could not reverse the sign of the frequency mismatch required to maintain a persistent *V*-leading-*D* state after T22 entrainment (**Figure 4D**).

Near the selected operating point, a ±5% change in period-memory strength altered the phase gap by 0.056 h on average, compared with 0.075 h for coupling-memory strength. The corresponding mean changes in dorsal period were 0.025 and 0.016 h (**Table S3**). These local responses were therefore similar and did not independently distinguish the mechanisms. The more informative distinction was the global response structure: period adaptation changed *Δω* continuously while preserving *K*_eff_, whereas coupling adaptation modified the interaction itself and could alter the existence and stability of the *D*-*V* phase relationship. Within the constrained coupling-only comparator tested here, intrinsic-period adaptation provided a more coherent common control axis across the five lighting schedules. This result does not exclude more general coupling-adaptation mechanisms in which additional coupling terms are allowed to vary.

### Bifurcation analysis and predictions for untested schedules

Bifurcation analysis showed that the availability and stability of *D*-*V* phase-locked states depended on both the schedule-specific frequency mismatch and *D*-to-*V* coupling (**Figure 5A**). The photoperiod conditions retained equilibrium branches over a broad range around the fitted *K_VD_* value, whereas the T-cycle conditions crossed no-lock regions. These curves emphasize that the same fitted coupling can support different phase organizations because each light history establishes a different early-DD *τ_D_* and therefore a different *Δω*.

Without further parameter adjustment, the model predicted distinct *in vivo* and *ex vivo* responses under 18:6 LD, T23, and T25 (**Figure 5B–D**; **Table 3**). The predicted *in vivo* behavioral periods were 23.579 h after 18:6 LD, 23.497 h after T23, and 24.086 h after T25. Removal of feedback from *X* produced *ex vivo D*/*V* periods of 23.333/23.497 h after 18:6 LD, 24.260/24.260 h after T23, and 23.172/23.333 h after T25. The corresponding *D*-*V* gaps were 1.088, −0.527, and 1.297 h. Thus, the model predicted a stronger long-photoperiod response under 18:6 LD, an intermediate short-period behavioral aftereffect after T23, and an intermediate long-period aftereffect after T25, while preserving an *in vivo*/*ex vivo* dissociation under the T-cycles.

**Table 3.** Model predictions for untested photoperiod and T-cycle schedules.

| Condition | Readout | *In vivo* (h) | *Ex vivo* (h) |
| --- | --- | --- | --- |
| 18:6 LD | Behavioral period | 23.579 | — |
| | $D$ period | | 23.333 |
| | $V$ period | | 23.497 |
| | $D$-$V$ phase gap | — | 1.088 |
| T23 | Behavioral period | 23.497 | — |
| | $D$ period | | 24.260 |
| | $V$ period | | 24.260 |
| | $D$-$V$ phase gap | — | -0.527 |
| T25 | Behavioral period | 24.086 | — |
| | $D$ period | | 23.172 |
| | $V$ period | | 23.333 |
| | $D$-$V$ phase gap | — | 1.297 |

**Notes.** Predictions were generated with the selected parameter set and no additional fitting. Positive $D$ - $V$ phase gaps indicate that $D$ leads $V$ ; a negative gap indicates that $V$ leads $D$ . *Ex vivo* behavioral and $X$ period readouts were not interpreted.

## Discussion

The present study integrates published behavioral and *ex vivo* SCN measurements with a simple *D*-*V*-*X* oscillator model to address why circadian aftereffects appear concordant under photoperiod manipulation but become inverted after non-24-h T-cycle entrainment. The model reproduced the expected photoperiodic ordering of free-running period, with long photoperiod producing a shorter period and short photoperiod producing a longer period, while also reproducing the opposite ordering of behavioral and explanted SCN periods after T22 and T26. Importantly, this dissociation emerged when feedback from the systemic oscillator *X* was removed at explantation. The central implication is therefore not that either behavior or the explanted SCN provides an incorrect measure of circadian period. Rather, they represent two different dynamical configurations: an intact *in vivo* network in which the SCN interacts with extra-SCN rhythmic signals, and an isolated SCN network that reorganizes after those interactions are removed.

The behavioral-SCN period inversion observed after T-cycle entrainment has now been reported across several experiments. Behavioral period generally changes in the direction of the entraining T-cycle, whereas *Per1-luc* or PER2::LUC rhythms recorded from subsequently explanted SCN can change in the opposite direction. This pattern was observed after short and long T-cycles by Aton et al. (2004), Molyneux et al. (2008), and Azzi et al. (2017). The interaction between T-cycles and photoperiods reported by Tackenberg et al. (2020) further showed that T-cycle length and photoperiod influence partly separable properties of the circadian system: behavioral activity duration and *ex vivo* SCN phase dispersion were more consistently associated than behavioral and *ex vivo* SCN free-running periods. Thus, the inversion cannot be dismissed simply as experimental noise, but neither does it imply that all properties of the *in vivo* and explanted SCN are oppositely organized. Instead, period, phase angle, phase dispersion, and behavioral waveform appear to preserve different components of prior light exposure.

Reporter identity may contribute to this heterogeneity, although the present model places greater emphasis on the change in network context caused by explantation. Different molecular reporters are not interchangeable projections of a single uniform SCN oscillator. Myung et al. (2012) found that *Bmal1-ELuc* rhythms preserved period-related spatial organization that was less apparent in PER2::LUC recordings. Ono et al. (2017) subsequently demonstrated *in vivo* that *Per1-luc* and *Bmal1-Eluc* rhythms shifted at different rates after light exposure and were associated with different behavioral landmarks, with *Per1* more closely following activity onset and *Bmal1* changing gradually with activity offset. Simultaneous dual-reporter recordings have also shown that *Bmal1* and *Per2* rhythms can develop different periods within the same cultured SCN slice (Nishide et al., 2018). Reporter selection can therefore alter which component of the network dominates the measured ensemble rhythm. Nevertheless, reporter specificity alone does not fully explain why a rhythm recorded *in vivo* can differ from that expressed by the same tissue after isolation. Reporter identity is more appropriately considered a measurement modifier acting in combination with the altered network state produced by explantation.

The *D*-V architecture used here is supported by extensive evidence that photoperiod is represented through structured phase relationships within the SCN rather than through a homogeneous shift of every cellular oscillator. Long photoperiod broadens or divides the phase distribution of SCN neurons, and anatomically distinct groups can become separated in time. Evans et al. (2013) showed that core and shell SCN neurons can peak 6–12 h apart after long-day exposure and subsequently resynchronize through phase-dependent interactions involving both VIP and GABA_A_ signaling. Myung et al. (2015) linked seasonal phase separation to spatial differences in intracellular [Cl^-^] and to attractive and repulsive GABAergic interactions. Rohr et al. (2019) further showed that seasonal plasticity in GABA_A_ signaling is required for the recovery of SCN synchrony. Causal evidence for a light-responsive ventral region comes from optogenetic activation of VIP neurons, which is sufficient to increase SCN phase dispersion and shorten subsequent *ex vivo* SCN period in a manner resembling long-photoperiod exposure (Tackenberg et al., 2021). These findings make a light-responsive ventral oscillator coupled to a more indirectly entrained dorsal oscillator a biologically plausible coarse-grained description of SCN organization.

The model further assigns persistent schedule-dependent period plasticity primarily to the dorsal oscillator. This assumption is consistent with evidence that dorsal AVP-expressing cells contribute to both period regulation and network coherence. Selective deletion of *Bmal1* from AVP neurons markedly lengthens behavioral free-running period and disrupts dorsal SCN rhythmicity, indicating that clocks in this population are not passive outputs but contribute to the period and coupling properties of the SCN network (Mieda et al., 2015). Persistent changes in intrinsic oscillator state are also biologically plausible because T-cycle exposure produces durable transcriptional and DNA-methylation changes within the SCN, while photoperiod alters rhythmic expression of genes associated with light responses, neuropeptide signaling, ion channels, and GABAergic function (Azzi et al., 2014, 2017; Cox et al., 2024; Myung et al., 2015). The adaptive dorsal period in the present model should therefore be interpreted as a reduced representation of several possible molecular and cellular memory processes rather than as evidence that a single transporter or clock gene alone stores the aftereffect. In particular, chloride-dependent GABA_A_ plasticity provides a plausible connection between photic history, phase organization, and period, for which KCC2/NKCC1 could be one molecular mechanism of *τ_D_* adaptation (Myung et al., 2015; Rohr et al., 2019).

Our comparison of adaptive-period and adaptive-coupling formulations favored persistent intrinsic-period adaptation for reproducing the full set of period and phase organization results across photoperiod and T-cycle conditions. This should not be interpreted as evidence that coupling plasticity is unimportant. Experimental and theoretical studies show that coupling can alter ensemble period, entrainment range, phase dispersion, and the stability of SCN organization. Synaptic plasticity can generate behavioral aftereffects in oscillator-network models, and the phase distribution of cellular oscillators can itself change ensemble period (Beersma et al., 2017; Foster et al., 2019). In T22-desynchronized rats, mutual coupling between ventrolateral and dorsomedial SCN oscillators contributes directly to the free-running period observed after release into DD (Schwartz et al., 2022). Kuramoto-model analyses also indicate that effective coupling is weaker under long photoperiod and stronger under short photoperiod in young SCN networks (van Beurden et al., 2023). The distinction suggested by our results is therefore functional rather than absolute. In the reduced model, intrinsic frequency mismatch provides the persistent directional bias required to produce shorter or longer aftereffects across all schedules, whereas coupling determines whether that bias is transferred coherently between dorsal and ventral oscillators. Coupling also determines the size and stability of their phase separation and nonstationary dynamics after release into darkness. This division of roles is compatible with the experimentally observed state dependence of SCN coupling: GABA can promote communication between widely separated oscillator groups yet oppose synchrony when the network is already coherent (Evans et al., 2013). A model in which coupling alone carries all photoperiod and T-cycle memories consequently becomes highly sensitive near changes in dynamical regime, whereas adaptive intrinsic period provides a smoother control variable and coupling stabilizes the resulting phase organization.

An alternative explanation for population-level effect also exists. Macroscopic period depends jointly on coupling strength and the width of the intrinsic period distribution (Myung et al., 2025b), so changes we model as adaptation of a single dorsal period could in part reflect a change in dorsal period heterogeneity. The two are not distinguishable from the period measurements used here.

The systemic oscillator *X* offers a possible explanation for why the same light history can produce different periods *in vivo* and after SCN explantation. *X* should not be interpreted as a second unidentified master pacemaker with a single anatomical location. It is a lumped representation of rhythmic extra-SCN feedback that remains available to the SCN in the intact animal but is removed from an isolated slice. Classical transplantation studies established that the donor SCN strongly determines the baseline period of restored locomotor behavior (Ralph et al., 1990). However, subsequent transplantation experiments found that properties retained by the host, including prior T-cycle aftereffects, could transiently influence rhythms restored by the graft (Matsumoto et al., 1996). Furthermore, *Per1*-mutant mice can exhibit nearly normal behavioral rhythms despite weak or irregular rhythms in isolated SCN tissue, leading to the proposal that *in vivo* clock interactions compensate for abnormalities expressed in culture (Pendergast et al., 2009). Together, these findings do not establish the existence of *X*, but they challenge the common assumption that the period measured in an isolated SCN slice must necessarily match the circadian period governing behavior in the intact animal.

Within our framework, T22 and T26 expose an interaction that is less evident under ordinary photoperiod manipulation. In the fitted model, T26 shifts the dorsal SCN toward a relatively short-period, phase-advanced state. In the intact animal, the longer-period influence represented by *X* opposes this SCN tendency and produces a longer behavioral aftereffect. Removal of *X* at explantation reveals the shorter-period SCN state. T22 produces the complementary configuration: the dorsal oscillator becomes relatively slow and phase delayed, while a shorter-period systemic influence maintains the short behavioral aftereffect *in vivo*. The isolated SCN therefore expresses a longer period. Under photoperiod manipulations, the intrinsic SCN and systemic biases are not opposed in the same manner, allowing behavioral and explant periods to retain the same ordering. This provides a unified explanation for why photoperiod and T-cycle aftereffects can involve overlapping SCN mechanisms while generating different behavioral-explant relationships. It is also consistent with the broader experimental conclusion that T-cycle length and light duration are not encoded as a single interchangeable variable (Tackenberg et al., 2020).

The intermediate T23 and T25 conditions, and longer photoperiod 18:6 LD are especially useful because they provide out-of-sample tests rather than additional fitting targets. The model predicts schedule-specific changes in behavioral period, *ex vivo* dorsal and ventral period, and the early-DD *D*-*V* phase gap. Measurements should therefore include not only a single free-running period but also the first several days of regional phase dynamics. A coupling-dominated mechanism would be expected to produce greater nonlinearity, instability, or between-slice variability near a transition between dynamical regimes. In contrast, a primarily intrinsic-period mechanism predicts more continuous changes in regional period, with coupling determining how rapidly the dorsal and ventral oscillators converge. Measuring only the final ensemble period would obscure this distinction.

Several limitations could temper the mechanistic interpretation of our model. First, dorsal and ventral variables represent aggregated oscillator populations and do not capture the full spatial and neurochemical diversity of the SCN. Second, *X* is phenomenological and its anatomical or physiological components remain unspecified. Third, the model treats schedule-induced period memory using a small number of state variables, whereas biological memory is likely distributed across transcriptional, epigenetic, ionic, synaptic, and circuit processes. Fourth, the literature synthesis combines studies differing in reporter, strain, age, sex, slice orientation, entrainment duration, and period-estimation window. These factors may contribute to apparent differences between studies. Fifth, behavioral, *in vivo* SCN, and *ex vivo* SCN periods were generally not measured longitudinally in the same animal. Sixth, although the model captured the principal ordering of periods and phase gaps, the magnitude of the T22 *D*-*V* separation remained underestimated, suggesting that additional spatial structure, nonlinear coupling, or heterogeneous cellular periods may be required. Finally, the fitted parameter set should not be interpreted as uniquely identified. Several parameter combinations may produce similar period and phase-gap outputs, and the inferred parameters of our model represent effective dynamical quantities rather than direct estimates of biological coupling strengths.

## Conclusion

In conclusion, the present results support a framework in which light history is encoded through adaptive changes in SCN dynamics, while the period expressed *in vivo* also depends on interactions that are lost during explantation. In our model, adaptive intrinsic period provides a parsimonious mechanism for representing prior light exposure, whereas *D*-*V* coupling shapes the stability and phase organization of the SCN network. The systemic oscillator *X* extends this framework without replacing the central role of the SCN, suggesting that behavioral timing emerges from the SCN interacting with the broader circadian system. This interpretation reframes the behavioral-explant discrepancy as a testable consequence of network disconnection rather than an inconsistency between measurements.

## Data and code availability

All data used in this study were obtained from previously published studies. The curated dataset containing the extracted behavioral and SCN period measurements, together with the corresponding study, reporter, lighting condition, and experimental metadata, is provided as Supplementary Data. Digitized source values, model-fitting targets, selected parameter sets, simulation outputs, and analysis code are available from the project repository:

https://github.com/vuonghtruong/Light-adaptive-SCN-two-oscillator-model

## Author contributions

V.H.T.: Conceptualization, methodology, software, formal analysis, investigation, data curation, visualization, and writing—original draft. J.M.: Conceptualization, supervision, funding acquisition, and writing—review and editing.

## Acknowledgements

We thank Cheng-en Tsai for preliminary exploratory simulations during the early development of the model. This work was supported by the National Science and Technology Council (NSTC) (114-2320-B-038-052-MY3), and by the Higher Education Sprout Project from the Ministry of Education (MOE), Taiwan. V.H.T. is the recipient of the NSTC PhD Dissertation Writing Award for Doctoral Candidates in the Humanities and Social Sciences (115-2424-H-038-001-DR).

## Declaration of generative AI use

During preparation of this manuscript, the authors used OpenAI Codex to assist with code cleanup, refactoring, debugging, and improvements to code readability. The tool was not used to generate experimental data, formulate the scientific hypothesis, select the model architecture, interpret the results, or draw the conclusions. All AI-assisted code was reviewed, tested, and validated by the authors, who take full responsibility for the final code and manuscript.

## References

Albus, H., Bonnefont, X., Chaves, I., Yasui, A., Doczy, J., van der Horst, G. T. J., & Meijer, J. H. (2002). Cryptochrome-deficient mice lack circadian electrical activity in the suprachiasmatic nuclei. Current Biology, 12(13), 1130–1133. 10.1016/s0960-9822(02)00923-5

Aton, S. J., Block, G. D., Tei, H., Yamazaki, S., & Herzog, E. D. (2004). Plasticity of Circadian Behavior and the Suprachiasmatic Nucleus Following Exposure to Non-24-Hour Light Cycles. Journal of Biological Rhythms, 19(3), 198–207. 10.1177/0748730404264156

Azzi, A., Dallmann, R., Casserly, A., Rehrauer, H., Patrignani, A., Maier, B., Kramer, A., & Brown, S. A. (2014). Circadian behavior is light-reprogrammed by plastic DNA methylation. Nature Neuroscience, 17(3), 377–382. 10.1038/nn.3651

Azzi, A., Evans, J. A., Leise, T., Myung, J., Takumi, T., Davidson, A. J., & Brown, S. A. (2017). Network dynamics mediate circadian clock plasticity. Neuron, 93(2), 441–450. 10.1016/j.neuron.2016.12.022

Beersma, D. G. M., Gargar, K. A., & Daan, S. (2017). Plasticity in the Period of the Circadian Pacemaker Induced by Phase Dispersion of Its Constituent Cellular Clocks. Journal of Biological Rhythms, 32(3), 237–245. 10.1177/0748730417706581

Buijink, M. R., Olde Engberink, A. H. O., Wit, C. B., Almog, A., Meijer, J. H., Rohling, J. H. T., & Michel, S. (2020). Aging Affects the Capacity of Photoperiodic Adaptation Downstream from the Central Molecular Clock. Journal of Biological Rhythms, 35(2), 167–179. 10.1177/0748730419900867

Campuzano, A., Vilaplana, J., Cambras, T., & Díez-Noguera, A. (1998). Dissociation of the rat motor activity rhythm under T cycles shorter than 24 hours. Physiology & Behavior, 63(2), 171–176. 10.1016/s0031-9384(97)00416-2

Ciarleglio, C. M., Axley, J. C., Strauss, B. R., Gamble, K. L., & McMahon, D. G. (2011). Perinatal photoperiod imprints the circadian clock. Nature Neuroscience, 14(1), 25–27. 10.1038/nn.2699

Cox, O. H., Gianonni-Guzmán, M. A., Cartailler, J.-P., Cottam, M. A., & McMahon, D. G. (2024). Transcriptomic Plasticity of the Circadian Clock in Response to Photoperiod: A Study in Male Melatonin-Competent Mice. Journal of Biological Rhythms, 39(5), 423–439. 10.1177/07487304241265439

Evans, J. A., Leise, T. L., Castanon-Cervantes, O., & Davidson, A. J. (2013). Dynamic interactions mediated by non-redundant signaling mechanisms couple circadian clock neurons. Neuron, 80(4), 10.1016/j.neuron.2013.08.022

Fernandez DC, Chang YT, Hattar S, and Chen SK (2016) Architecture of retinal projections to the central circadian pacemaker. Proc Natl Acad Sci USA, 113, 6047–6052.

Foster, S., Christiansen, T., & Antle, M. C. (2019). Modeling the Influence of Synaptic Plasticity on After-effects. Journal of Biological Rhythms, 34(6), 645–657. 10.1177/0748730419871189

Herzog, E. D., Takahashi, J. S., & Block, G. D. (1998). Clock controls circadian period in isolated suprachiasmatic nucleus neurons. Nature Neuroscience, 1(8), 708– 713. 10.1038/3708

Honma, S., Shirakawa, T., Katsuno, Y., Namihira, M., & Honma, K. (1998). Circadian periods of single suprachiasmatic neurons in rats. Neuroscience Letters, 250(3), 157–160. 10.1016/s0304-3940(98)00464-9

Keene, N. E., Pourmir, F., Hang, E., Ozturget, M., McCreedy, D. A., & Jones, J. R. (2026). A direct SCN-to-DMH output pathway organizes circadian behavioral timing (p. 2026.05.21.726959). bioRxiv. 10.64898/2026.05.21.726959

Kitaguchi, Y., Komori, S., Tei, H., & Uriu, K. (2025). Attractive and repulsive couplings between circadian pacemaker neurons promote entrainment to light-dark cycles with after-effect (p. 2025.11.20.689452). bioRxiv. 10.1101/2025.11.20.689452

Liu, C., Weaver, D. R., Strogatz, S. H., & Reppert, S. M. (1997). Cellular construction of a circadian clock: Period determination in the suprachiasmatic nuclei. Cell, 91(6), 855–860. 10.1016/s0092-8674(00)80473-0

Matsumoto, S., Basil, J., Jetton, A. E., Lehman, M. N., & Bittman, E. L. (1996). Regulation of the phase and period of circadian rhythms restored by suprachiasmatic transplants. Journal of Biological Rhythms, 11(2), 145–162. 10.1177/074873049601100207

Mieda, M., Ono, D., Hasegawa, E., Okamoto, H., Honma, K.-I., Honma, S., & Sakurai, T. (2015). Cellular clocks in AVP neurons of the SCN are critical for interneuronal coupling regulating circadian behavior rhythm. Neuron, 85(5), 1103–1116. 10.1016/j.neuron.2015.02.005

Molyneux, P. C., Dahlgren, M. K., & Harrington, M. E. (2008). Circadian entrainment aftereffects in suprachiasmatic nuclei and peripheral tissues *in vitro*. Brain Research, 1228, 127–134. 10.1016/j.brainres.2008.05.091

Moore, R. Y., Speh, J. C., & Leak, R. K. (2002). Suprachiasmatic nucleus organization. Cell and Tissue Research, 309(1), 89–98. 10.1007/s00441-002-0575-2

Myung, J., Hong, S., DeWoskin, D., De Schutter, E., Forger, D. B., & Takumi, T. (2015). GABA-mediated repulsive coupling between circadian clock neurons in the SCN encodes seasonal time. Proceedings of the National Academy of Sciences, 112(29), E3920–E3929. 10.1073/pnas.1421200112

Myung, J., Hong, S., Hatanaka, F., Nakajima, Y., De Schutter, E., & Takumi, T. (2012). Period coding of Bmal1 oscillators in the suprachiasmatic nucleus. The Journal of Neuroscience: The Official Journal of the Society for Neuroscience, 32(26), 8900–8918. 10.1523/JNEUROSCI.5586-11.2012

Myung, J., & Pauls, S. D. (2018). Encoding seasonal information in a two-oscillator model of the multi-oscillator circadian clock. The European Journal of Neuroscience, 48(8), 2718–2727. 10.1111/ejn.13697

Myung, J., Vitet, H., Truong, V. H., & Ananthasubramaniam, B. (2025a). The role of the multiplicity of circadian clocks in mammalian systems. Sleep Medicine, 131, 106518. 10.1016/j.sleep.2025.106518

Myung, J., Vitet, H., & Tiong, S. Y. X. (2025b). Modeling the geometry of circadian synchronization and period across aging. Biogerontology, 26(4), 157. 10.1007/s10522-025-10303-1

Nakamura, W., Honma, S., Shirakawa, T., & Honma, K. (2002). Clock mutation lengthens the circadian period without damping rhythms in individual SCN neurons. Nature Neuroscience, 5(5), 399–400. 10.1038/nn843

Nishide, S., Honma, S., & Honma, K. (2018). Two coupled circadian oscillations regulate Bmal1-ELuc and Per2-SLR2 expression in the mouse suprachiasmatic nucleus. Scientific Reports, 8, 14765. 10.1038/s41598-018-32516-w

Ono, D., Honma, S., Nakajima, Y., Kuroda, S., Enoki, R., & Honma, K.-I. (2017). Dissociation of Per1 and Bmal1 circadian rhythms in the suprachiasmatic nucleus in parallel with behavioral outputs. Proceedings of the National Academy of Sciences, 114(18), E3699–E3708. 10.1073/pnas.1613374114

Patke, A., Young, M. W., & Axelrod, S. (2020). Molecular mechanisms and physiological importance of circadian rhythms. Nature Reviews. Molecular Cell Biology, 21(2), 67–84. 10.1038/s41580-019-0179-2

Pendergast, J. S., Friday, R. C., & Yamazaki, S. (2009). Endogenous Rhythms in Period1 Mutant Suprachiasmatic Nuclei *In Vitro* Do Not Represent Circadian Behavior. The Journal of Neuroscience, 29(46), 14681–14686. 10.1523/JNEUROSCI.3261-09.2009

Ralph, M. R., Foster, R. G., Davis, F. C., & Menaker, M. (1990). Transplanted suprachiasmatic nucleus determines circadian period. Science, 247(4945), 975– 978. 10.1126/science.2305266

Rohr, K. E., Pancholi, H., Haider, S., Karow, C., Modert, D., Raddatz, N. J., & Evans, J. (2019). Seasonal plasticity in GABA_A_ signaling is necessary for restoring phase synchrony in the master circadian clock network. eLife, 8, e49578. 10.7554/eLife.49578

Schwartz, M. D., Cambras, T., Díez-Noguera, A., Campuzano, A., Oda, G. A., Yamazaki, S., & de la Iglesia, H. O. (2022). Coupling Between Subregional Oscillators Within the Suprachiasmatic Nucleus Determines Free-Running Period in the Rat. Journal of Biological Rhythms, 37(6), 620–630. 10.1177/07487304221126074

Starnes, A. N., & Jones, J. R. (2023). Inputs and Outputs of the Mammalian Circadian Clock. Biology, 12(4), 508. 10.3390/biology12040508

Tackenberg, M. C., Hughey, J. J., & McMahon, D. G. (2020). Distinct Components of Photoperiodic Light Are Differentially Encoded by the Mammalian Circadian Clock. Journal of Biological Rhythms, 35(4), 353–367. 10.1177/0748730420929217

Tackenberg, M. C., Hughey, J. J., & McMahon, D. G. (2021). Optogenetic stimulation of VIPergic SCN neurons induces photoperiodic-like changes in the mammalian circadian clock. The European Journal of Neuroscience, 54(9), 7063–7071. 10.1111/ejn.15442

van Beurden, A. W., Meylahn, J. M., Achterhof, S., Buijink, R., Olde Engberink, A., Michel, S., Meijer, J. H., & Rohling, J. H. T. (2023). Reduced Plasticity in Coupling Strength in the Aging SCN Clock as Revealed by Kuramoto Modeling. Journal of Biological Rhythms, 38(5), 461–475. 10.1177/07487304231175191

Vitaterna, M. H., Takahashi, J. S., & Turek, F. W. (2001). Overview of Circadian Rhythms. Alcohol Research & Health, 25(2), 85–93.

Welsh, D. K., Logothetis, D. E., Meister, M., & Reppert, S. M. (1995). Individual neurons dissociated from rat suprachiasmatic nucleus express independently phased circadian firing rhythms. Neuron, 14(4), 697–706. 10.1016/0896-6273(95)90214-7

Yamaguchi, Y., Suzuki, T., Mizoro, Y., Kori, H., Okada, K., Chen, Y., Fustin, J.-M., Yamazaki, F., Mizuguchi, N., Zhang, J., Dong, X., Tsujimoto, G., Okuno, Y., Doi, M., & Okamura, H. (2013). Mice genetically deficient in vasopressin V1a and V1b receptors are resistant to jet lag. Science, 342(6154), 85–90. 10.1126/science.1238599

Yamazaki, S., Straume, M., Tei, H., Sakaki, Y., Menaker, M., & Block, G. D. (2002). Effects of aging on central and peripheral mammalian clocks. Proceedings of the National Academy of Sciences of the United States of America, 99(16), 10801–10806. 10.1073/pnas.152318499

