## Supplementary Information for "A two-oscillator SCN model with period adaptation and systemic feedback captures photoperiod and T-cycle aftereffects *in vivo* and in explants"

### Analytical driver-based sensitivity of the reduced model

Sensitivity was expressed in terms of the two direct drivers of the reduced phase-gap equation, $\Delta\omega$ and $K_{\mathrm{eff}}$, rather than raw derivatives with respect to $\tau_{D,0}$ and $K_{VD}$. For a locked solution,

$$\phi^* = \sin^{-1}\left(\frac{\Delta\omega}{K_{\mathrm{eff}}}\right), \qquad r = \frac{\Delta\omega}{K_{\mathrm{eff}}}$$

$$d\phi^* = \frac{dr}{\sqrt{1-r^2}}, \qquad \frac{dr}{r} = \frac{d\Delta\omega}{\Delta\omega} - \frac{dK_{\mathrm{eff}}}{K_{\mathrm{eff}}}$$

Thus, equal fractional changes in $\Delta\omega$ and $K_{\mathrm{eff}}$ have equal magnitude but opposite effects on the equilibrium phase in the reduced locked model. With $\tau_V$ and $K_{DV}$ fixed, their biological parameter mappings are:

$$\frac{\partial\Delta\omega}{\partial\tau_D} = -\frac{2\pi}{\tau_D^2}, \qquad \frac{\partial K_{\mathrm{eff}}}{\partial K_{VD}} = 1$$

The corresponding phase-gap derivatives are:

$$\frac{\partial\phi^*}{\partial\Delta\omega} = \frac{1}{\sqrt{K_{\mathrm{eff}}^2 - \Delta\omega^2}}$$

$$\frac{\partial\phi^*}{\partial K_{\mathrm{eff}}} = -\frac{\Delta\omega}{K_{\mathrm{eff}}\sqrt{K_{\mathrm{eff}}^2 - \Delta\omega^2}}$$

$$\frac{\partial\phi^*}{\partial K_{\mathrm{eff}}} = -r\,\frac{\partial\phi^*}{\partial\Delta\omega}$$

Within the stable principal branch, changing $K_{\mathrm{eff}}$ has relatively little effect on the equilibrium phase gap when |r| is small, whereas changing $\Delta\omega$ continues to shift the equilibrium. Sensitivity to either driver increases as |r| approaches 1 and diverges at the fixed-point boundary. When $|r| > 1$, the reduced time-independent system has no stationary $D$-V phase relationship. These expressions assume $K_{\mathrm{eff}} > 0$; for $K_{\mathrm{eff}} < 0$, stability occurs on the alternative equilibrium branch.

The common locked angular frequency is

$$\Omega = \omega_V + K_{VD}\frac{\Delta\omega}{K_{\mathrm{eff}}}$$

Holding the coupling constants fixed while varying $\Delta\omega$ gives

$$\frac{\partial\tau_{\mathrm{lock}}}{\partial\Delta\omega} = -\frac{2\pi}{\Omega^2}\frac{K_{VD}}{K_{\mathrm{eff}}}, \qquad \tau_{\mathrm{lock}} = \frac{2\pi}{\Omega}$$

whereas varying $K_{VD}$ with $K_{DV}$ and $\Delta\omega$ fixed gives

$$\frac{\partial\tau_{\mathrm{lock}}}{\partial K_{\mathrm{eff}}} = \frac{2\pi}{\Omega^2}\frac{K_{DV}\Delta\omega}{K_{\mathrm{eff}}^2}$$

Their relationship is

$$\frac{\partial\tau_{\mathrm{lock}}}{\partial K_{\mathrm{eff}}} = -\frac{K_{DV}}{K_{VD}}\, r\, \frac{\partial\tau_{\mathrm{lock}}}{\partial\Delta\omega}$$

When the frequency mismatch is small relative to coupling, changing $K_{\mathrm{eff}}$ has little effect on the stationary period. This helps explain why the curves in **Figure 4F** are comparatively flat over much of their range, whereas changing $\Delta\omega$ produces a clearer and more continuous period response in **Figure 4E**.

**Table S1. Selected model parameters and optimization scores.**

| Category | Parameter | Value |
|---|---|---|
| Intrinsic periods | $\tau_{D,0}$ | 23.9592 h |
| | $\tau_V$ | 24.0329 h |
| | $\tau_{X,0}$ | 23.4241 h |
| Adaptive-period gains | $g_D$ | 7.99447 |
| | $g_X$ | 33.6108 |
| Memory gains | $g_{\Phi,D}$ | –1.89013 |
| | $g_{P,D}$ | 0.0582106 |
| | $g_{\Phi,X}$ | 1.91262 |
| | $g_{P,X}$ | –0.378657 |
| $D$-$V$ coupling | $K_{VD}$ | 0.016000 rad $h^{-1}$ |
| | $K_{DV}$ | 0.012425 rad $h^{-1}$ |
| Systemic coupling | $K_{XD}$ | 0.398597 rad $h^{-1}$ |
| | $K_{XV}$ | 0.197781 rad $h^{-1}$ |
| | $K_{SX}$ | 0.365419 rad $h^{-1}$ |
| Light input | $A_L$ | 0.339446 |
| | $b_V$ | 18.3508 |
| Optimization metrics | Raw screening objective | $7.29852 \times 10^6$ |
| | Diagnostic-compromise score | 14.6647 |
| | Lomb-Scargle selection score | 14.6647 |

**Notes.** $D$, $V$, and $X$ denote the dorsal SCN, ventral SCN, and systemic oscillator, respectively. The raw screening objective used the computationally inexpensive phase-slope surrogate and is not directly comparable with the Lomb-Scargle final-selection score. Running several Astropy Lomb-Scargle periodograms for every candidate would make the search substantially slower and is difficult to include efficiently inside the parallel Numba kernel. For screening, period is approximated from the unwrapped phase:

$$\widehat{\omega} = \frac{\sum_t (t-\bar{t})[\phi(t)-\bar{\phi}]}{\sum_t (t-\bar{t})^2},\ \hat{\tau}_{\text{slope}} = \frac{2\pi}{\widehat{\omega}}.$$

**Table S2. Reduced early-DD phase-locking diagnostics at the fitted operating point.**

| Condition | $\Delta\omega$ (rad $h^{-1}$) | $K_{eff}$ (rad $h^{-1}$) | $\Delta\omega/K_{eff}$ | $R=\|\Delta\omega/K_{eff}\|$ | Reduced fixed point exists |
|---|---|---|---|---|---|
| SP | 0.000196 | 0.003575 | 0.055 | 0.055 | Yes |
| LP | 0.002821 | 0.003575 | 0.789 | 0.789 | Yes |
| LD | 0.001500 | 0.003575 | 0.420 | 0.420 | Yes |
| T22 | –0.004296 | 0.003575 | –1.202 | 1.202 | No |
| T26 | 0.007324 | 0.003575 | 2.049 | 2.049 | No |

**Notes.** The diagnostic uses the condition-specific mean early-DD $\tau_D$ and the fitted $K_{eff}$. $R \leq 1$ is necessary for a fixed point in the reduced time-independent $D$-$V$ system. The full model remains time-dependent because the memory states continue to relax in DD.

**Table S3. Full-model sensitivity to adaptive-period and adaptive-coupling gain scales.**

| Architecture | Gain change | Mean \|Δgap\| (h) | Max \|Δgap\| (h) | Mean \|ΔD period\| (h) | Mean \|ΔV period\| (h) | Lock to no lock | No lock to lock |
|---|---|---|---|---|---|---|---|
| Adaptive period | ±5% | 0.056 | 0.133 | 0.025 | 0.017 | 0/10 | 0/10 |
| Adaptive period | ±50% | 0.494 | 1.133 | 0.269 | 0.230 | 1/10 | 1/10 |
| Adaptive period | ±100% | 1.098 | 2.692 | 0.480 | 0.442 | 1/10 | 2/10 |
| Adaptive coupling | ±5% | 0.075 | 0.516 | 0.016 | 0.026 | 0/10 | 0/10 |
| Adaptive coupling | ±50% | 0.379 | 1.986 | 0.099 | 0.095 | 0/10 | 2/10 |
| Adaptive coupling | ±100% | 0.470 | 1.984 | 0.182 | 0.095 | 0/10 | 2/10 |

**Notes.** Lock transitions were classified relative to the selected gain scale $s = 1$. 'Lock to no lock' denotes loss of a reduced fixed point after perturbation; 'no lock to lock' denotes acquisition of a reduced fixed point. Each magnitude contains ten schedule-direction comparisons: five schedules multiplied by the positive and negative gain perturbations.

**Table S4. Numerical and reduced analytical sensitivities at the selected adaptive gains.**

| Condition | Architecture | *R* | Numerical $d\phi/d$driver (h) | Reduced $d\phi/d$driver (h) | Numerical $d\tau/d$driver ($h^2$ $rad^{-1}$) | Reduced $d\tau/d$driver ($h^2$ $rad^{-1}$) |
|---|---|---|---|---|---|---|
| SP | Adaptive period | 0.055 | 79.665 | 280.139 | −0.000 | −408.654 |
| SP | Adaptive coupling | 0.123 | −2.056 | −18.972 | 0.000 | 21.126 |
| LP | Adaptive period | 0.789 | 80.149 | 455.300 | −163.521 | −374.377 |
| LP | Adaptive coupling | 0.149 | −8.896 | — | 46.705 | — |
| LD | Adaptive period | 0.420 | 79.745 | 308.142 | 0.000 | −391.068 |
| LD | Adaptive coupling | 1.856 | −4.106 | — | 135.023 | — |
| T22 | Adaptive period | 1.202 | 79.863 | — | −155.268 | — |
| T22 | Adaptive coupling | 0.029 | −0.691 | −1.041 | 0.000 | 1.178 |
| T26 | Adaptive period | 2.049 | 82.843 | — | −93.996 | — |
| T26 | Adaptive coupling | 0.032 | 29.588 | — | −44.936 | — |

**Notes.** Numerical derivatives are central differences from the full time-dependent simulation and are normalized by the schedule-specific change in the corresponding reduced model. Reduced analytical derivatives are reported only where the relevant principal locked branch exists; a dash indicates that the derivative is undefined under the stated criterion. Analytical and numerical values need not coincide because the full simulation includes memory decay, transient dynamics, phase wrapping, and Lomb-Scargle period estimation.
